# First Comprehensive Examination of the Molecular Phylogenetics of Saki Monkeys, Genus *Pithecia* Desmarest, 1804, Reveals an Unexpectedly Low Taxonomic Diversity

**DOI:** 10.1101/2025.07.06.663372

**Authors:** Jean P. Boubli, Felipe E. Silva, Pedro Senna Bittencourt, Romina Batista, Rodrigo Costa-Araújo, Cleuton Miranda, Rogerio Rossi, Marcelo Gordo, Paris Badrock, Russell Mittermeier, João Valsecchi, Maria N. F. Silva, Mariluce Messias, Anneke H. van Heteren, Christian Roos, Wilsea Figueiredo, Anthony B. Rylands, Tomas Hrbek, Izeni Farias

## Abstract

Saki monkeys (*Pithecia*) are found exclusively in the Amazon, ranging from the Guiana Shield in the east to the Andean foothills in the west. The taxonomy of this genus is complex due to indeterminate type localities, numerous synonyms, sexual dichromatism and considerable geographical diversity in coat colour patterns. Previous assessments have indicated two to four species and a variable few subspecies based on morphological analyses of museum specimens. Just two species have been consistently recognised—*Pithecia pithecia* and *Pithecia monachus*. This study presents the first comprehensive molecular phylogeny of the genus, using mitochondrial cytochrome *b* sequences from 137 individuals across their range and a ddRADseq genomic analysis on 34 individuals. The phylogenetic results revealed three main clades: (1) Guiana Shield sakis, (2) Sedimentary Basin and Brazilian Shield sakis, and (3) a basal *P. albicans* south of the Rio Solimões. The ddRADseq data further clarified species boundaries and relationships, identifying six distinct species: *P. chrysocephala* and *P. pithecia* sensu Hershkovitz (1987) on the Guiana Shield; *P. albicans* in central Amazonia south of the Rio Solimões; *P. monachus* in western Amazonia; *P. irrorata* between the rios Purus, Tapajós, and Juruena; and *P. vanzolinii* in a limited area east of the Rio Juruá.

## Text Introduction

The saki monkeys (*Pithecia*) occur exclusively in Amazonia, from the Guiana Shield in the east to the Andean foothills in the west but are absent from the west bank of the Rio Branco to the north bank of the Rio Japurá-Caquetá (Hershkovitz 1987, Boubli et al., 2015) (Fig. 1). Sakis are sympatric with bearded sakis (*Chiropotes*) in a large part of their ranges north of the lower Rio Amazonas and with bald uakaris (*Cacajao*) in central and western Amazonia (Boubli et al., 2015). Perhaps to avoid competition with bearded sakis, which are canopy dwellers, the sakis are mostly observed using the lower and middle strata of the forest (Ayres, 1989). Weighing from about 1.5 kg (*P. pithecia*) to 3 kg (*P. albicans*), sakis are the smallest of the pitheciine primates and have locomotor adaptations for vertical clinging and leaping, which is unusual in the Platyrrhini but does appear in the Callitrichidae (marmosets, Goeldi’s monkey, tamarins and lion tamarins) and is also a characteristic of the Indriidae (sifakas, simponas and woolly lemurs) and the Tarsiidae. They are amongst the least studied of the Neotropical primates, not only because they are found mostly in remote places but also because studying them in the field has proven challenging due to their behaviour of moving quietly and freezing when perceiving danger or human activity (Pinto et al., 2013).

**Figure 1.**
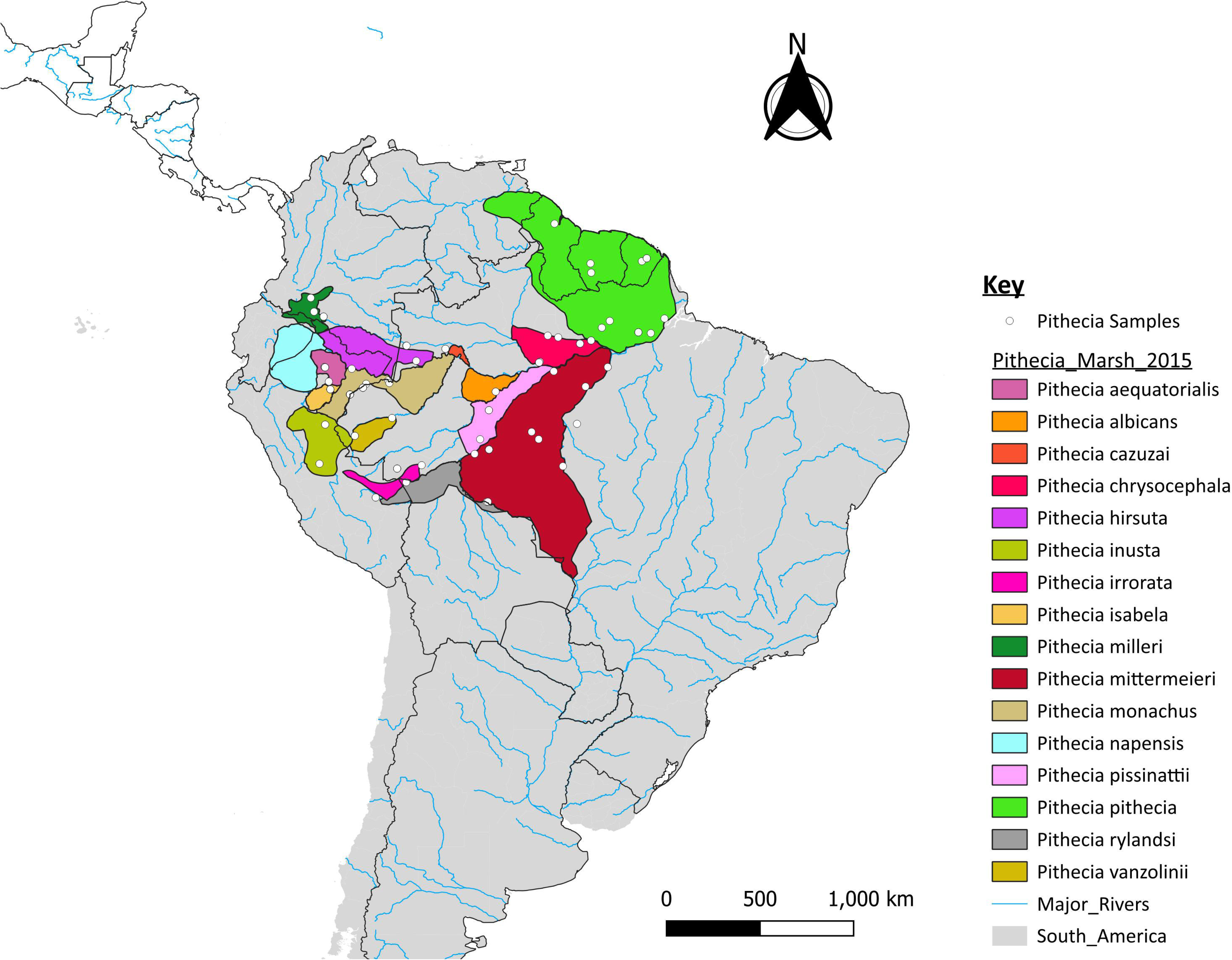
Geographic distribution of 16 *Pithecia* species according to Marsh (2014). White dots: Localities of the samples used in the molecular analysis of this study.

The taxonomy of *Pithecia* has a complex history, largely due to the lack of preserved type specimens and precise type localities (Hershkovitz, 1987; Marsh, 2014). Moreover, some species present marked sexual dichromatism, which has confounded taxonomic identification and assessments. Cruz Lima (1945) listed five species: *Pithecia pithecia* (Linnaeus, 1766), *P. chrysocephala* I. Geoffroy Saint-Hilaire, 1850, *P. monachus* (É. Geoffroy Saint-Hilaire, 1812), *P. milleri* J. A. Allen 1914, and *P. albicans* Gray, 1860. Cabrera (1957) listed just two species, *P. pithecia* and *P. monachus*, with a subspecies, *P. m. capillamentosa* Spix, 1823. Hill (1960) listed those of Cruz Lima but with *chrysocephala* as a subspecies of *P. pithecia*, and *capillamentosa, milleri* and *albicans* as subspecies of *monachus.* Napier (1976) was the most conservative, recognising only *P. pithecia* and *P. monachus*.

Hershkovitz (1979, 1987) carried out the first comprehensive taxonomic revision of the genus in more recent years and proposed five species and eight taxa. A second taxonomic assessment of the genus was carried out by Marsh (2014) who, following the Phylogenetic Species Concept, increased the number of species to 16, which included Hershkovitz’s eight taxa, revalidated three species, and described five ‘new’ species. Both taxonomic proposals were based very largely on pelage patterns. Currently, the IUCN’s Primate Specialist Group recognizes 15 saki species, following Marsh (2014), except in the recognition of *Pithecia rylandsi* Marsh, 2014, considered to be a junior synonym of the, evidently sympatric, *Pithecia irrorata* Gray, 1843 (see Rylands et al., 2024; also Rylands and Mittermeier, 2024 http://www.primate-sg.org/primates_of_neotropics).

Figueiredo (2005) conducted the sole molecular phylogenetic study of *Pithecia*, based on sequences of the mitochondrial cytochrome *b* gene (cyt *b*) of museum specimens from across the distribution of the genus. Her phylogenetic analysis revealed three well-supported clades: one comprising sakis from the Guiana Shield (*P. pithecia* and *P. chrysocephala*, *sensu* Hershkovitz, 1987 and Marsh, 2014), a second from the Amazonian Sedimentary Basin and Brazilian Shield (*P. aequatorialis* Hershkovitz, 1987, *P. irrorata irrorata*, *P. irrorata vanzolinii* Hershkovitz, 1987, *P. monachus monachus, P. monachus milleri*, *sensu* Hershkovitz 1987), and a third of *P. albicans*, a species with a relatively small geographic range. These findings corroborated, to a large degree, the taxonomic hypothesis of Hershkovitz (1987).

To gain further insights into the taxonomy of this genus, we carried out a comprehensive molecular phylogenetic investigation of *Pithecia*. We sequenced the mitochondrial cytochrome *b* gene from fresh tissue samples and museum specimens, including the three syntypes of *P. hirsuta* (Spix, 1823). We also generated a genome-wide sampling (ddRADseq *sensu* Patterson, 2012) for a selected number of specimens. Based on our results, we re-evaluate the taxonomy of *Pithecia* as proposed by Hershkovitz (1987) and Marsh (2014).

## Materials and Methods

For our analysis (species assignments for our samples), we followed Marsh (2014), considering the 16 species (including *P. rylandsi*) and their respective geographic distributions as published in the IUCN Red List. They are: *P. monachu*s; *P. irrorata*; *P. vanzolinii*; *P. albicans*; *P. chrysocephala*; *P. pithecia*; *P. milleri*; *P. aequatorialis*; *P. hirsuta*; *P. inusta* Spix, 1823; *P. napensis* Lönnberg, 1938; *P. mittermeieri* Marsh, 2014; *P. rylandsi*; *P. pissinattii* Marsh, 2014; *P. isabela* Marsh, 2014; and *P. cazuzai* Marsh, 2014.

### Cytochrome b dataset

The samples for the cytochrome *b* gene included 14 of the species recognized by Marsh (2014). No samples were available for *P. napensis* and *P. rylandsi*. To employ a broad geographic sampling, we combined Figueiredo’s (2005) alignment, which consisted of 103 cytochrome *b* sequences from fresh tissue samples (N=20) and historical museum specimens (N=83) with 31 newly generated cytochrome *b* sequences from fresh tissue samples collected in the field, and the cytochrome *b* sequence from the three syntypes of *P. hirsuta*, also newly generated for this study. We also included sequences of *Cacajao melanocephalus* (N=1) and *Chiropotes sagulatus* (N=1) as outgroups.

Permits to conduct fieldwork and sample collections were provided by the Brazilian Ministry of the Environment agencies – Chico Mendes Institute for Biodiversity Conservation (ICMBio) (SISBio 42111-1, 42111-2, 42111-3, 8937-1) and the Brazilian Institute of Environment and Renewable Natural Resources (IBAMA) (License N° 005/2005 – CGFAU/LIC).

Details for DNA extraction and amplification are found in Figueiredo (2005). Briefly, DNA from fresh tissue was isolated following the phenol-chloroform protocol (Hillis et al., 1996). DNA isolation for historical samples was done in a clean lab after UV and DNAaway (E&K Scientific) treatment for half an hour. A 2 mm^2^ of the voucher’s skin was used as input. After three washes with sodium hypochlorite 5% (O’Rourke et al., 2000) followed by three washes with ddH_2_O, skins had their DNA isolated using QIAmp or DNAsy kits (Quiagen). A table with information about the primers used for cyt *b* amplification can be found in Supplementary Table 1. For the fresh samples collected in this study, we used the phenol-chloroform protocol for DNA extraction from Sambrook et al. (1989) and the protocol detailed in Silva et al. (2022) for DNA quantification and cyt *b* gene amplification and sequencing. The cyt *b* sequence of the three syntypes of *P. hirsuta* was extracted from complete mitochondrial genomes of these specimens, which were produced using a whole-genome shotgun sequencing approach described in Boubli et al. (2021). We used Geneious Pro 4.8.5 for the cyt *b* sequence editing and assembly and aligned sequences using the Mafft online server (Katoh and Standley, 2013; Katoh et al., 2017) under the iterative refinement option (FFT-NS-i Standard) (Katoh et al., 2002, 2017).

### Genomic dataset

We obtained good-quality DNA for genomic analysis for 34 of our fresh tissue samples (Table 1), which included *P. chrysocephala* (N=8), *P. pithecia* (N=7), *P. albicans* (N=2), *P. vanzolinii* (N=1), *P. hirsuta* (N=4), *P. mittermeieri* (N=9), and *P. pissinattii* (N=3). As outgroups, we included *Chiropotes israelita* (N=2). For genomic sequencing, we carried out a double digest restriction-site-associated DNA sequencing protocol (ddRADseq) adapted from Patterson et al. (2012) that allows a simultaneous digestion, ligation, and barcoded adapter incorporation (https://github.com/legalLab/protocols-scripts/tree/master/ddRAD (see for example Boubli et al., 2018; Collins and Hrbek, 2018; Silva et al., 2018). We assessed the quality of raw reads using FastQC v.0.11.8 (Andrews, 2018) and kept only those reads with PHRED scores >30 for subsequent analyses. We used the software pipeline iPYRAD (Eaton and Overcast, 2020) to process the reads and map them to the reference genome of *Pithecia pithecia* (GenBank: GCA_028551515.1). We set a minimum sample per locus to 2 (∼5% of samples) to build our catalogue of loci and retrieved 18,017 loci in a matrix with 5,302,691 positions (63.55% of missing data). We then used a sample map to include at least one individual per species in our dataset, and we ended up with 1,661 loci in a sequence matrix of 503,945 positions and 24.74% missing sites.

**Table 1.**
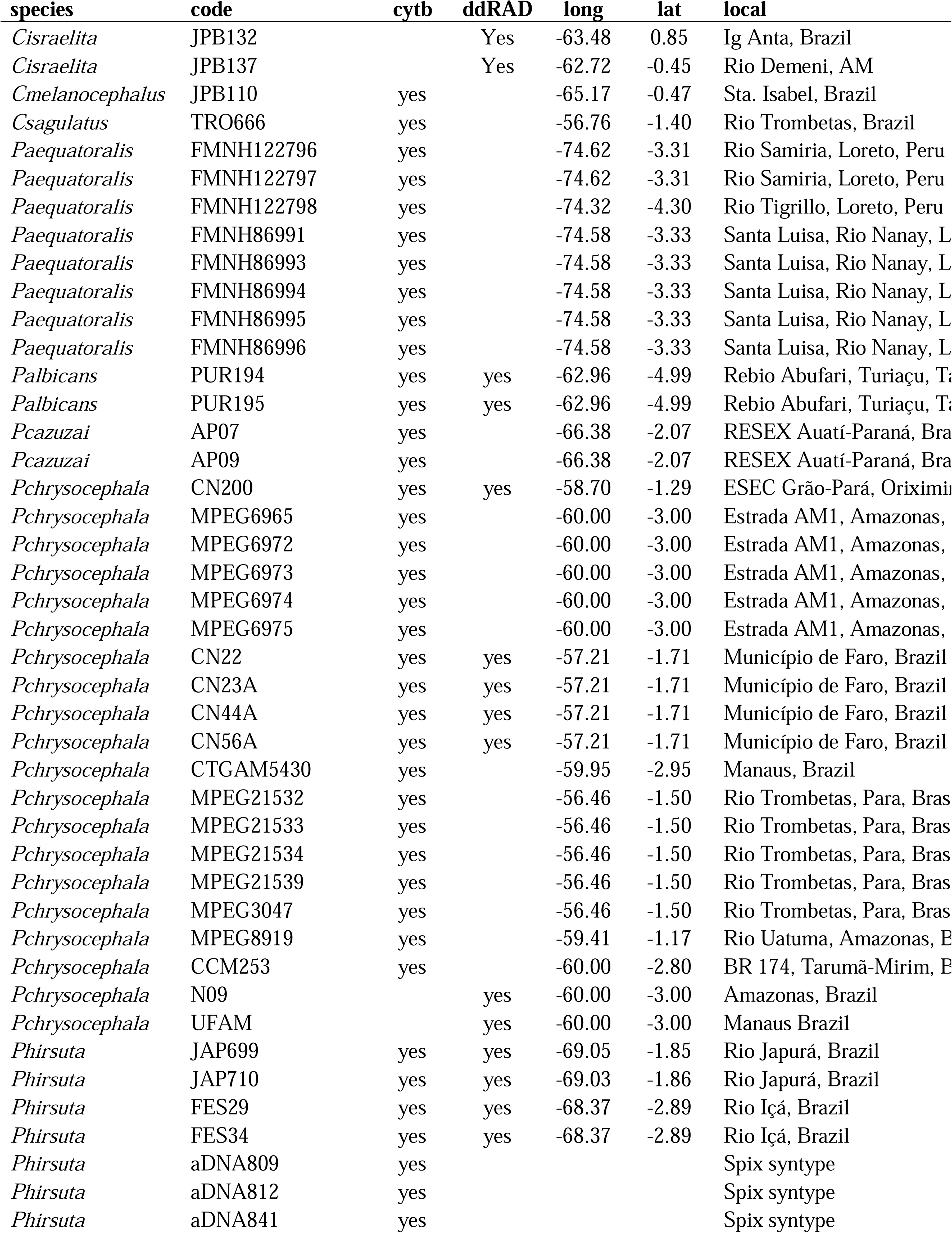
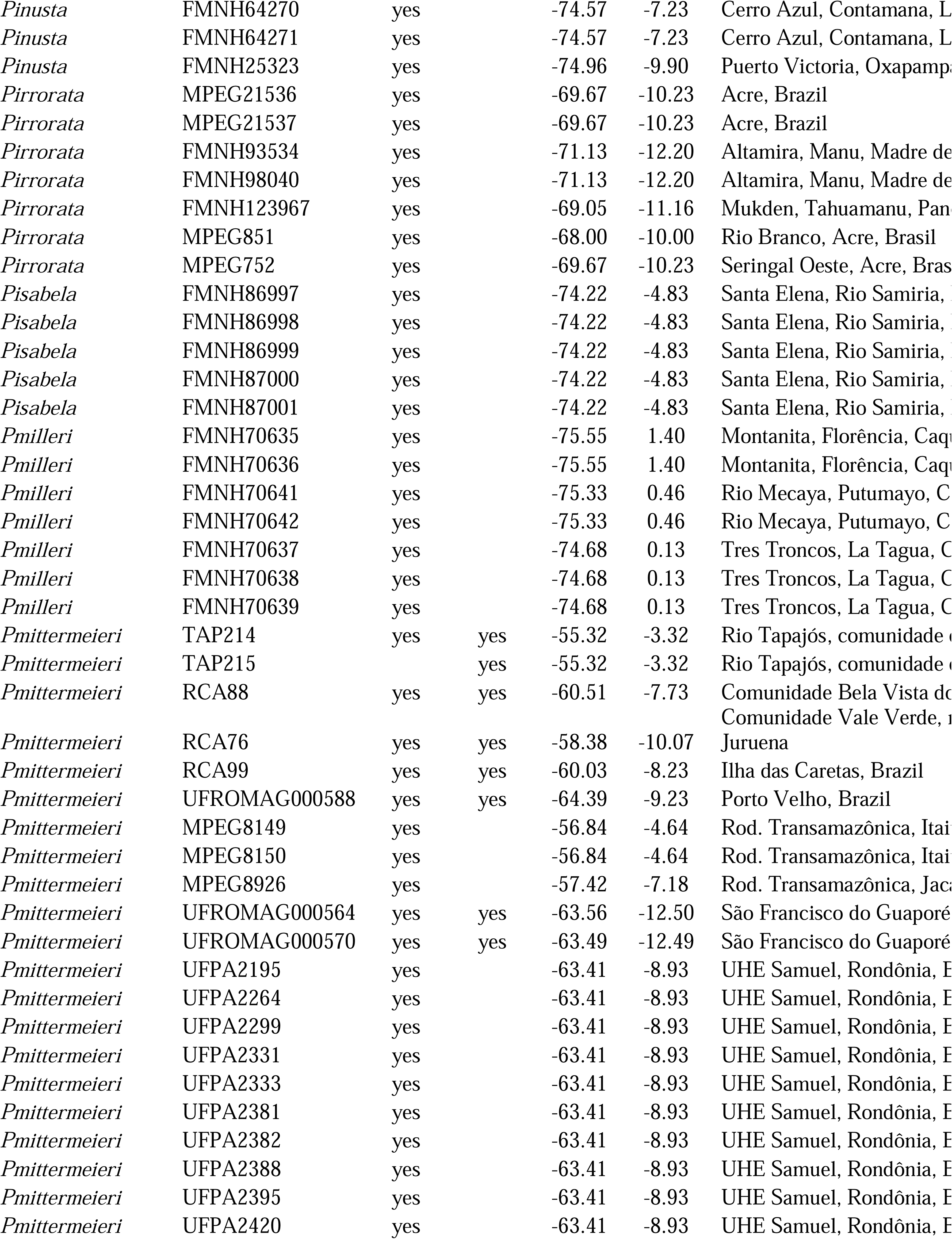
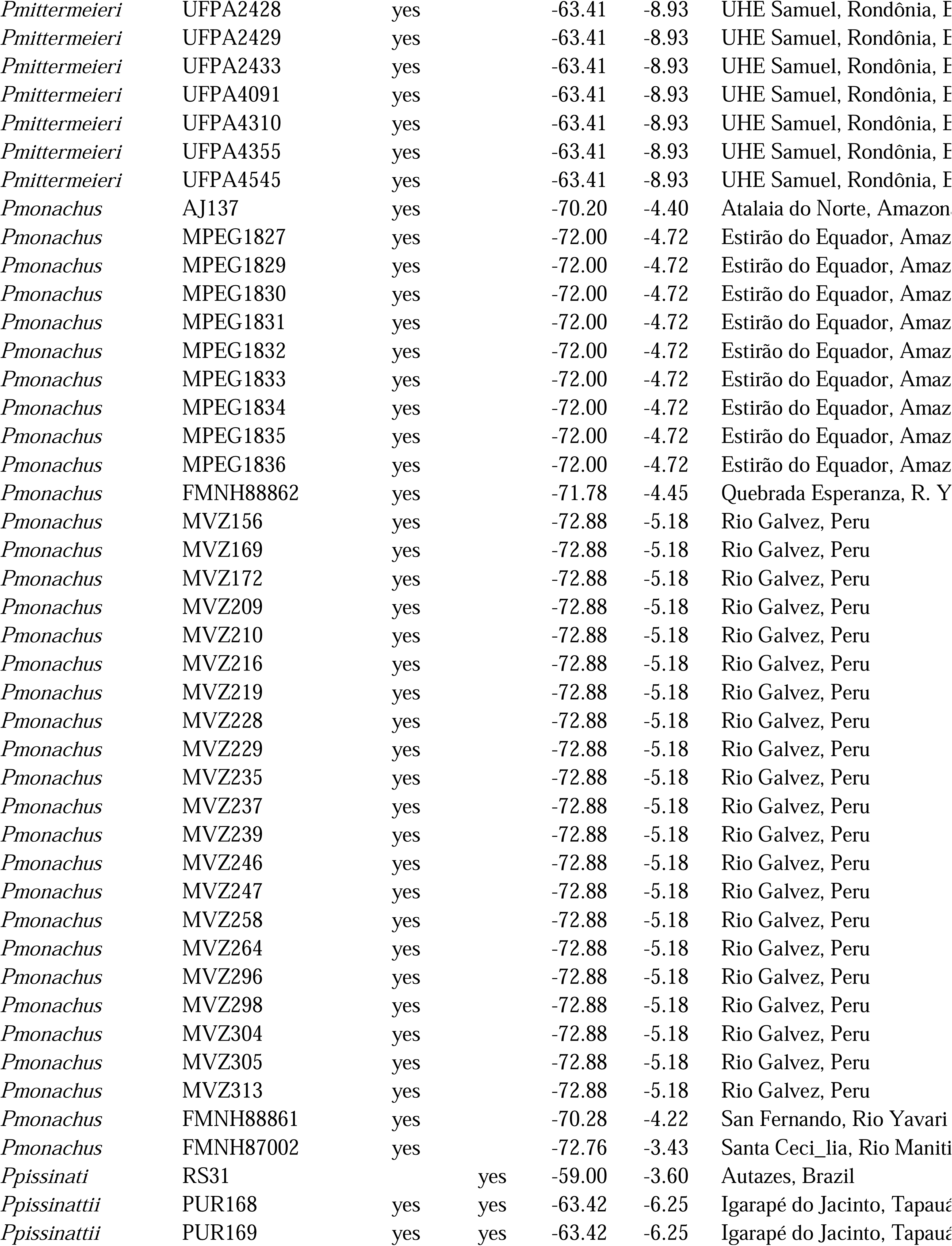
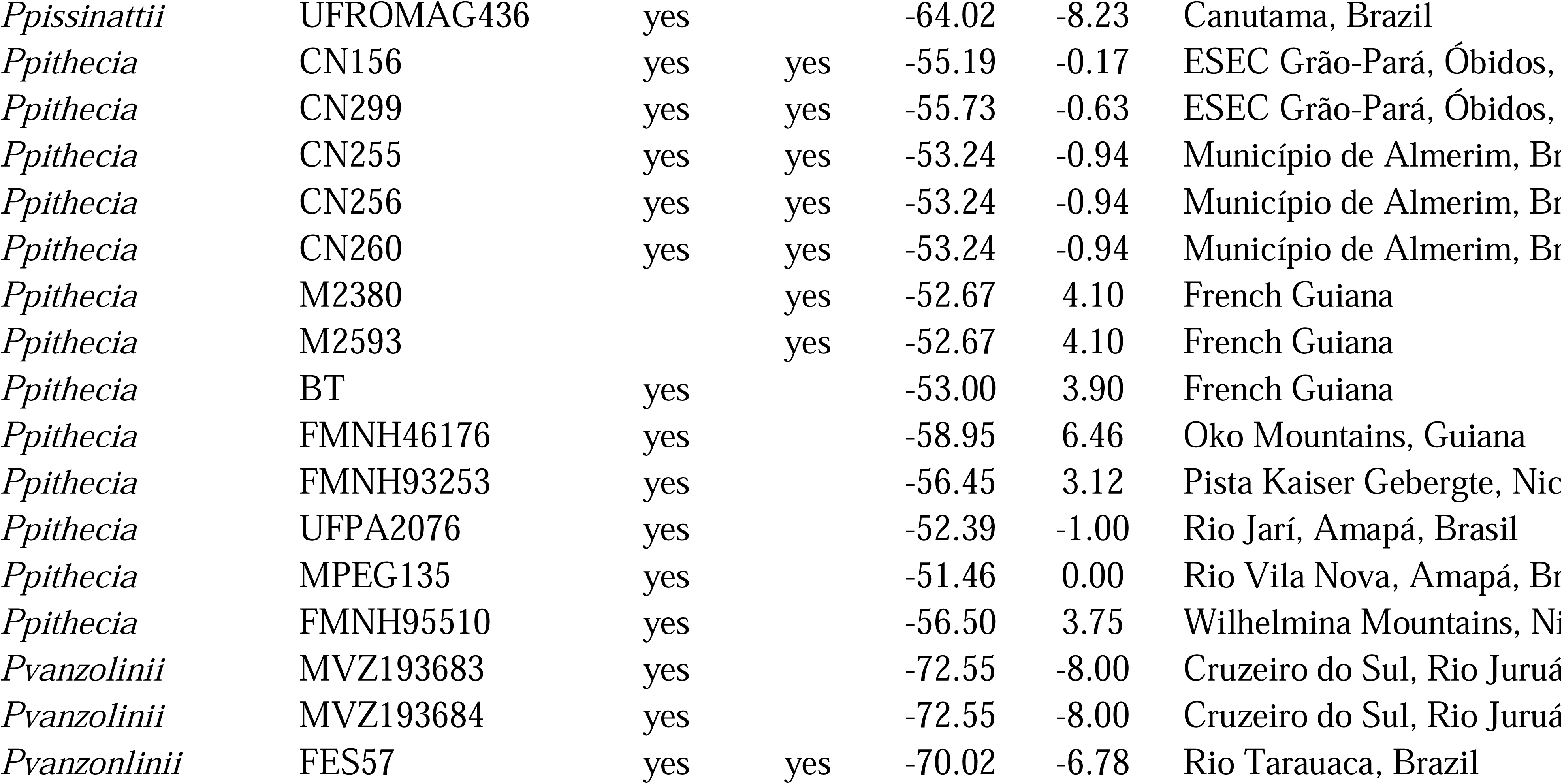
Pithecia samples used in this study.

### Phylogenetic analyses

Maximum-likelihood (ML) phylogenetic inferences were carried out using the cyt *b* sequences and the ddRAD dataset separately. The ML tree was generated with IQTree v.1.6.12 (Nguyen et al., 2015), using the ModelFinder algorithm (Kalyaanamoorthy et al., 2017) as implemented in the W-IQ-Tree (http://iqtree.cibiv.univie.ac.at; Trifinopoulos et al., 2016) to select the best substitution model, and the ultrafast bootstrap (UFBoot) (Hoang et al., 2018; Minh et al., 2020) with 1,000 pseudo-replicates to assess branch support.

### Species delimitation

For species delimitation analyses we used the maximum credibility Bayesian tree as input. To run our tree, we first reduced the total dataset, originally containing 137 cytochrome *b* sequences plus two outgroups, to a new dataset containing only 100 unique haplotypes using the function ‘hap_collapse’ from ‘delimtools’ R package (https://github.com/legalLab/delimtools) in R. Then we ran the Bayesian inference in BEAST v2.6.7 (Bouckaert et al., 2019) using the following settings: site model (TPM2u+I+G) as estimated by the BEAST2 package bModelTest 1.2.1 (Bouckaert and Drummond, 2017); strict molecular clock; coalescent constant population tree prior. We ran three independent runs with 20 million Markov Chain Monte Carlo (MCMC) generations, annotating trees and parameters every 2,000 generations. Convergence between chains was observed by checking the values of effective sample sizes (ESS > 200) and the stationarity of the chain using TRACER 1.7.1 (Rambaut et al., 2018). We combined trees and parameters and subsampled it at a frequency of 6,000 generations, removing the first 10% generations of each run as *burn-in* using LogCombiner (Drummond et al., 2012) to produce a dataset with 9,000 topologies which we used to search for the maximum credibility tree using TREEANNOTATOR (Bouckaert et al., 2019). This Bayesian tree was used as input for the following tree-based, single-locus species delimitation analyses: GMYC, the Generalized Mixed Yule Coalescent model (Pons et al., 2006; Fujisawa and Barraclough, 2013), implemented in the R package ‘splits 1.0-20’ (Ezard et al., 2021); bGMYC, a Bayesian implementation of GMYC in the R package ‘bGMYC 1.0.3’ (Reid & Carstens, 2012) with custom modifications for compatibility with R versions above 4.0.0; PTP, the Poisson Tree Process (Zhang et al., 2013); and mPTP, the multi-rate Poisson Tree Process (Kapli et al., 2016), both implemented on the ‘mptp 0.2.5’ software. For both PTP and mPTP, we estimated a ML tree using the maximum credibility tree as a start using the function ‘pml_bb’ of R package ‘phangorn’ 2.11.1 (Schliep, 2011). We also used the entire cyt *b* dataset as input for LocMin, the Local Minima method, which is a distance-based optimising and clustering method implemented in ‘spider 1.4-2’ (Brown et al., 2012). We then combined all species delimitation outputs by using the ‘delimtools’ function ‘delim_join’, calculated a majority-vote consensus by using the function ‘delim_consensus’ and visualised the haplotype tree alongside all species delimitation results and consensus using the function ‘delim_autoplot.’

### Genetic variation

Based on the main lineages recovered in the phylogenetic analyses with ddRAD data, we carried out a principal component analysis (PCA) using a matrix composed of single-nucleotide polymorphisms (SNP) generated in the iPYRAD pipeline. Accordingly, we first ran a PCA including *P. hirsuta*, *P. vanzolinii*, *P. albicans*, *P. mittermeieri*, and *P. pissinattii*. After filtering and keeping only the SNPs presented in at least 50% of all samples, we obtained a matrix of 10,217 SNPs, with 17.70% of missing data. To run the PCA, we used a subset of 2,812 unlinked SNPs. After that, we ran a second PCA, including only *P. chrysocephala* and *P. pithecia* data and obtained a matrix with 20,847 SNPs, of which 5,441 were unlinked (14.46% of missing data).

## Results

### Cyt b Dataset

The cyt *b* alignment contained 139 sequences (137 from *Pithecia* spp. and 2 from outgroups), with a length of 1,138 base pairs, 296 variable sites, and 190 parsimony-informative sites. In the phylogenetic inference, we retrieved three main lineages with high support in the ML inference: 1) *P. chrysocephala* and *P. pithecia* (from the Guiana Shield), 2) *P. albicans* (from the lower Rio Tefé), and 3) all remaining taxa (Fig. 2). The two currently recognised species from the Guiana Shield were recovered as paraphyletic. *Pithecia albicans* was retrieved as monophyletic. The syntypes of *P. hirsuta* grouped with modern samples of *P. hirsuta* from the Rio Japurá. Two samples of *P. monachus* (MPEG 1831, FMNH 88862) were recovered as sister to the clade containing all samples of species from the sedimentary Basin and Brazilian Shield. In the Bayesian inference, we observed some geographical structuring within the Sedimentary Basin and the Brazilian shield: 1) *P. pissinatti*, *P. mittermeieri*, and *P. irrorata* in the Purus-Madeira and Madeira-Tapajós interfluvia, 2) *P. vanzolinii*, *P. cazuzai*, *P. hirsuta*, *P. monachus*, and *P. milleri*, west of the rios Juruá and Japurá, and 3) *P. isabela*, *P. inusta*, and *P. aequatorialis* from the Andean foothills (Fig. 3).

**Figure 2.**
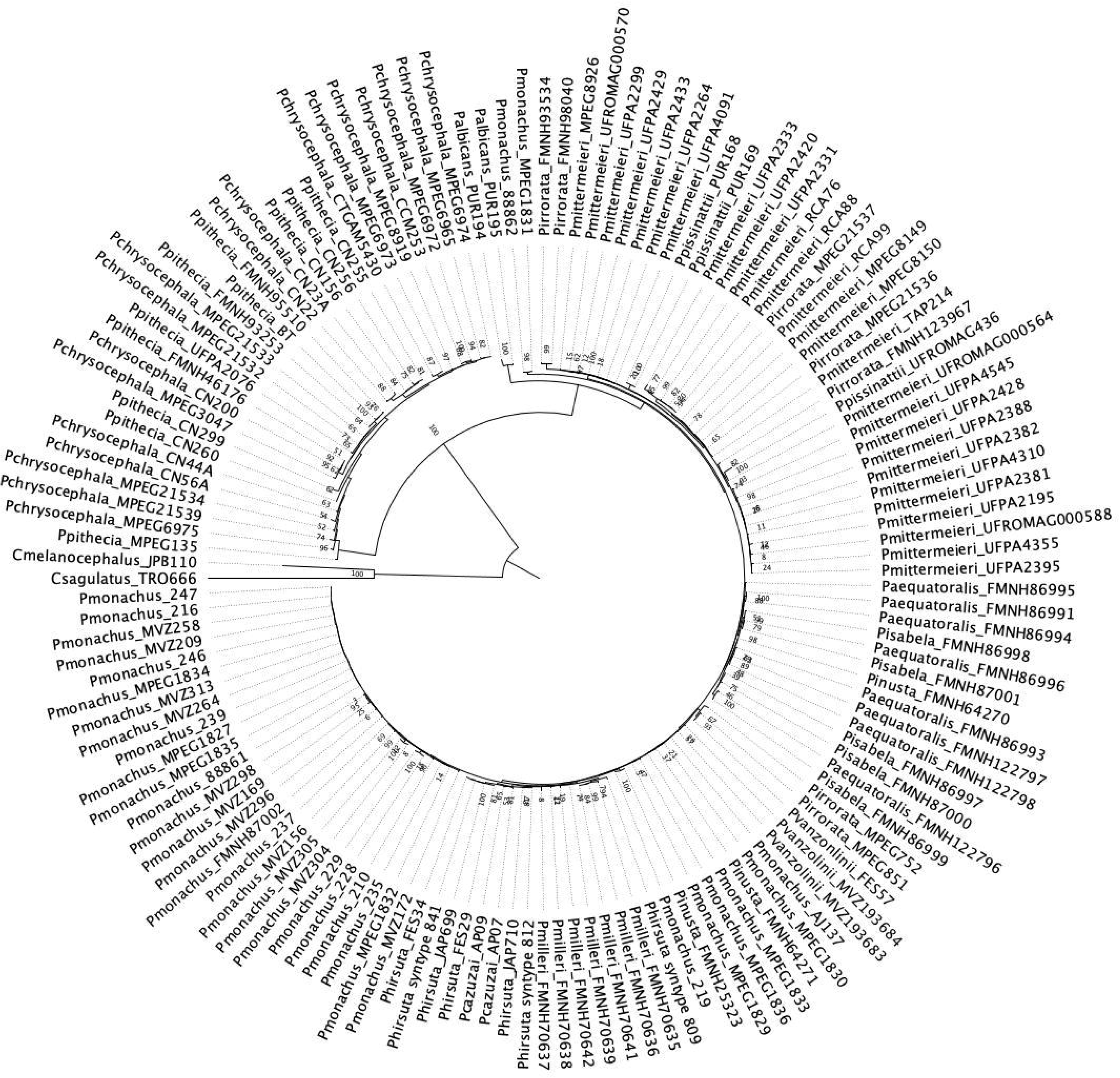
Maximum likelihood tree retrieved from the cyt *b* alignment of 137 Pithecia samples and two outgroups; numbers in the centre correspond to bootstrap values.

**Figure 3.**
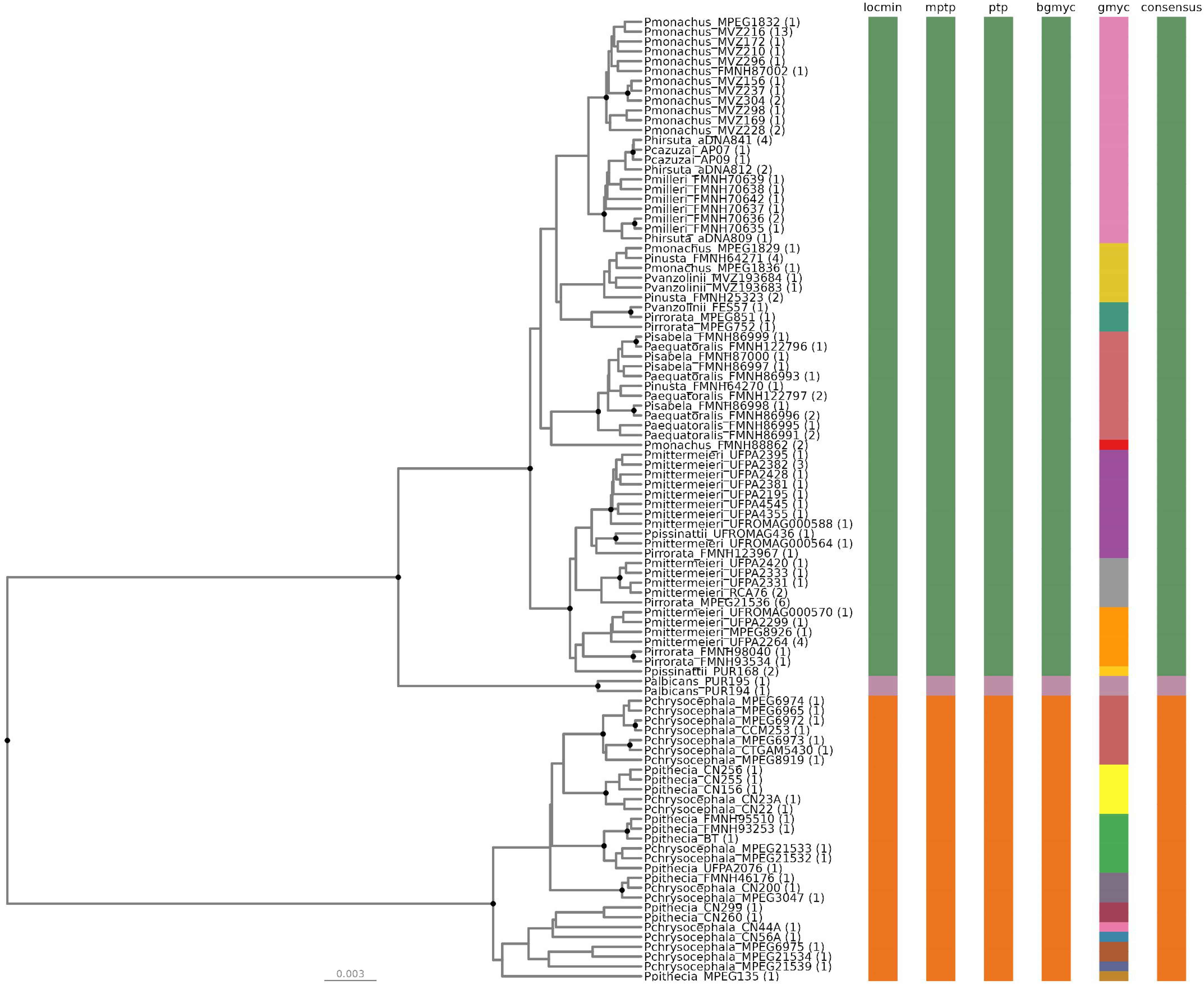
Maximum clade credibility tree from 9,000 posterior trees generated using BEAST 2.6.7. Dataset comprised 100 unique haplotypes (from a total of 137) of *Pithecia* cytochrome b sequences. Bayesian posterior probabilities above 0.95 are shown as dark nodes. Species delimitations are shown per method as colored bars where colors indicate different species partitions. Matching colors across different methods indicate congruence between methods for a given species partition. The Consensus bar was calculated by using the majority-vote method of ‘delim_consensus’. The number of collapsed individuals per haplotype is indicated in parentheses. The figure was created in R 4.4.1 using the package ‘delimtools’.

With the exception of the GMYC method, all other methods of species delimitation identified three lineages within our cyt *b* dataset. The first lineage was composed of 11 saki species, namely *P. aequatorialis*, *P. cazuzai*, *P. hirsuta*, *P. inusta*, *P. irrorata*, *P. isabela*, *P. milleri*, *P. mittermeieri*, *P. monachus*, *P. pissinattii*, and *P. vanzolini*. The second lineage was composed of *P. albicans*, sister clade to the first lineage. The third lineage was composed of *P. chrysocephala* and *P. pithecia*—it was the most basal in our phylogeny. The GMYC method delimited a total of 20 lineages, splitting the first and third consensus lineages into smaller species partitions, which may indicate some degree of population structure. The LocMin method indicates a genetic distance threshold of 1.9% as the transition point between intra and interspecific genetic distances between *Pithecia* species (Fig. 3).

### Genomic Dataset

In the ddRAD genomic analysis, the saki monkeys from the Guiana Shield, *P. chrysocephala* and *P. pithecia*, were recovered as reciprocally monophyletic sister clades with 100% support. *Pithecia albicans* is a basal taxon of the saki monkeys from the Sedimentary Basin and the Brazilian Shield. *Pithecia hirsuta* is a monophyletic clade and the only ddRAD sequence of *P. vanzolinii* in our phylogeny is an offshoot of the clade that includes *P. mittermeieri + P. pissinatii* (Fig. 4).

**Figure 4.**
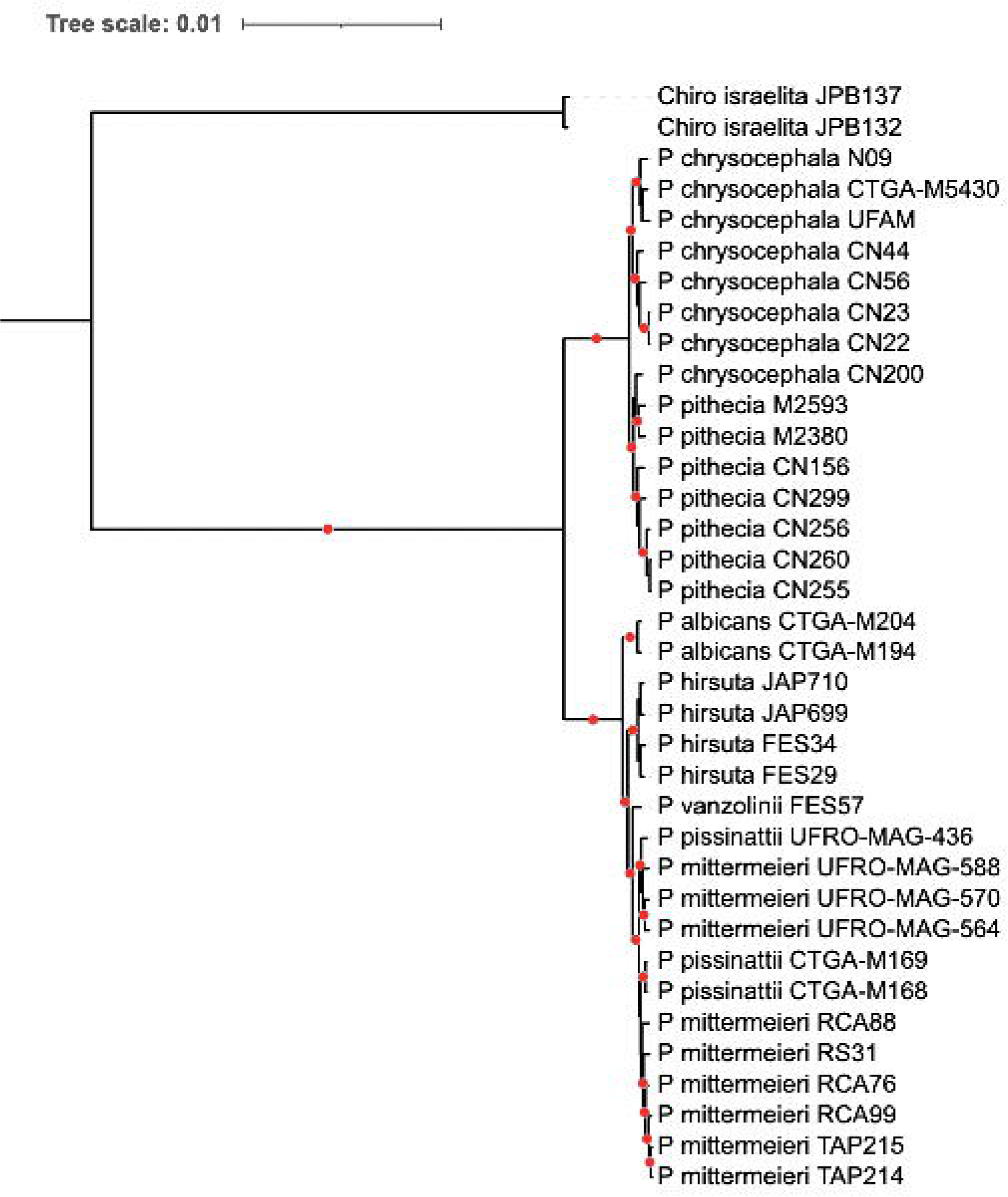
Maximum likelihood tree retrieved from the ddRAD concatenated matrix of loci showing the main *Pithecia* lineages. Red circles indicate nodes with 100% of bootstrap support.

In the PCA analysis, 42.1% of the genetic variation found in the Sedimentary Basin plus Brazilian Shield clade is explained by the first two components, with a clear distinction between *P. hirsuta*, *P. vanzolinii*, and *P. albicans*, and a cluster that includes both *P. pissinattii* and *P. mittermeieri* (Fig. 5A). In comparing components 1 and 3 *P. pissinattii* form an independent cluster from *P. mittermeieri* (Fig. 5B). For the Guiana Shield clade, the first two components explain 34.7% of the genetic variation, with *P. chrysocephala* and *P. pithecia* separated into two main clusters (Fig. 5C). Sample CN200 with predominant morphology of *P. pithecia sensu* Hershkovitz (1987) and Marsh (2014) appears isolated in the middle of the PCA charts (Figs. 5C and 5D).

**Figure 5.**
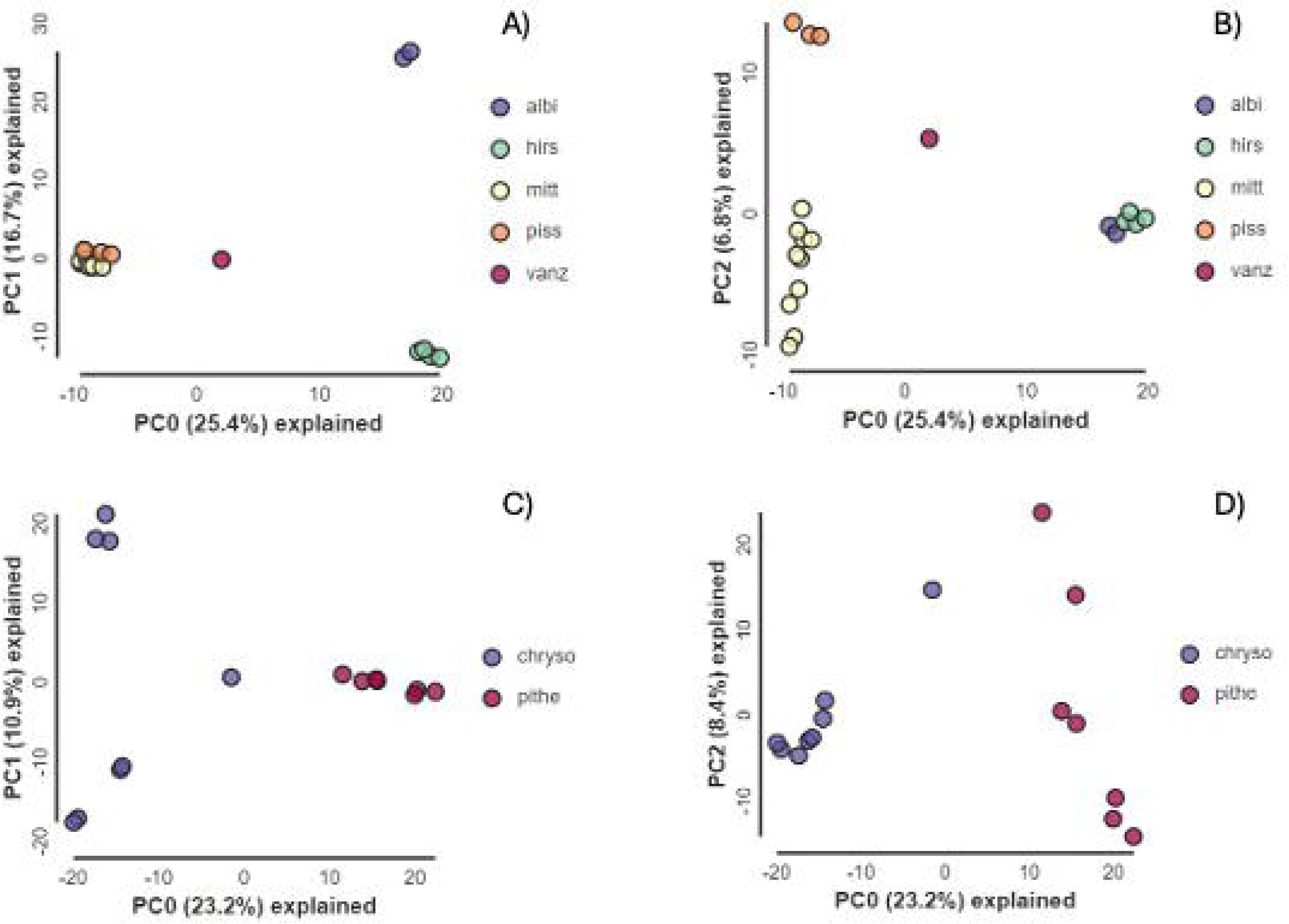
Principal component analysis (PCA) showing the *Pithecia* allele frequency variation using unlinked SNPs. A and B) The PCA using 2,812 unlinked SNPs for *Pithecia* from the Sedimentary basin and Brazilian shield. Most of the variation, explained by components 0 and 1, shows *P. mittermeieri* and *P. pissinattii* grouped in one single cluster, while *P. vanzolinii*, *P. hirsuta* and *P. albicans* are grouped into separated clusters each. C and D) The PCA using 5,441 unlinked SNPs for the Guiana shield sakis, shows *Pithecia chrysocephala* and *P. pithecia* separated into two main clusters, with the individual CN200 occupying an intermediary position between them (see text).

## Discussion

The two major recent taxonomic reviews of the sakis are those of Hershkovitz (1979, 1987) and Marsh (2014). Both were based on morphology and, principally, coat colour patterns, as well as the more or less evident sexual dichromatism prevalent in the genus. Hershkovitz (1987) recognised five species, three of them with subspecies, totalling eight taxa:

1. *Pithecia* group – representing both sakis from the Guiana Shield, north of the Rio Amazonas: *P. pithecia* (*P. pithecia pithecia*, *P. pithecia chrysocephala*), and
2. *Monachus* group – representing all sakis from the Sedimentary Basin and Brazilian Shield, south of the rios Amazonas-Solimões, Juruá and Napo: *P. albicans*, *P. aequatorialis* (first description), *P. irrorata* (*P. irrorata irrorata*, *P. irrorata vanzolinii* (first description), and *P. monachus* (*P. monachus monachus*, *P. monachus milleri*).

Marroig and Cheverud (2004; see also Marroig et al., 2003) carried out a study of the cranial morphometry of the taxa proposed by Hershkovitz (1987). They corroborated his taxonomic arrangement, identifying the two species groups—the diminutive *Pithecia pithecia* group and the rather larger *P. monachus* group*—*but argued that members of the *monachus* should all be considered species. They found *P. albicans* to be basal, followed by *P. aequatorialis*, and then the sister clades of [*P. vanzolinii*, *P irrorata*, *P. monachus*] and [*P. p. pithecia*, *P. p. chrysocephalus*]. Groves (2001, 2005), Norconk (2011) and Silva Júnior et al. (2013) followed the classification of Hershkovitz (1987).

Marsh (2014) recognised 16 taxa, all classified as species. They include the eight taxa recognised by Hershkovitz (1987), three forms that Hershkovitz considered synonyms of *monachus*, and five that were newly described:

1. *Pithecia inusta* from the basin of the Río Ucayali in Peru and the middle to upper Juruá in Brasil; *Pithecia napensis* from the Napo basin in Ecuador and Peru, the Río Santago in Ecuador, and the northern bank of the Río Marañón in Peru; *Pithecia hirsuta* from the north of the Río Napo in Peru, in the Colombian Trapezium north of the Río Amazonas to the Río Caquetá, and in the Içá-Putumayo basin in Peru and Brazil. Hershkovitz (1987) argued *hirsuta* was, along with *monachus* and *irrorata*, a name misapplied or misused throughout almost its entire history (p. 408).
2. The species newly described by Marsh (2014) were: *Pithecia isabela* in the Pacaya- Samiria basin in Peru; *Pithecia cazuzai* from between the mouth of the Rio Japurá to the Rio Solimões; *Pithecia mittermeieri* from the interfluvium of the rios Madeira, Tapajós, and Juruena; *Pithecia rylandsi* from extreme northwestern Bolivia and the Río Madre de Dios in Peru; and *Pithecia pissinattii* from between the lower rios Madeira and Purus in Brazil.

Marsh’s (2014) re-assessment of the taxonomy of this genus was carried out over 10 years and was exhaustive in its examination of 876 skins and 690 skulls kept in 36 museums in 28 cities in 17 countries—supplemented by hundreds of photographs. Notable is her exceptional capacity to discern differences in pelage colour and patterns related to the age and sex of the diverse specimens maintained in the collections, revealing, as a consequence the extraordinary geographic variation in phenotypes of a genus characterized by sexual dichromatism, with pelage colour and patterns changing (transitioning) as the sakis grow to adulthood. Marsh’s study showed that both features have confounded taxonomists attempting to unravel the taxonomy of this genus.

Despite this phenotypic variation, our study revealed an unexpectedly low phylogenetic diversity in the genus *Pithecia*, with six lineages in the ddRADseq phylogeny and three lineages in the cyt *b* phylogeny. Although our cyt *b* phylogeny analysis did not support *P. pithecia* and *P. chrysocephala* as separate species, as was proposed by Marroig and Cheverud (2004) and Marsh (2014), both the genomic and PCA analyses showed a clear separation between the two, with a clear geographic structure: *P. chrysocephala* restricted to the west and *P. pithecia* to the east of the Rio Trombetas, a right bank tributary of the Rio Amazonas. In the PCA, specimen CN200 that was collected on the left bank of the Rio Trombetas, and thus on the western limit of the range of *P. pithecia* (see Silva Junior et al., 2013) appeared isolated between *P. chrysocephala* and *P. pithecia*. It has a whitish face with orange extensions but with *P. pithecia* phenotype, *sensu* Hershkovitz (1987) and Marsh (2014), predominating (Cleuton Miranda, pers. obs.). Marsh (2014) suggested that samples from this region may be hybrids of the two, which could explain the placement of CN200 in our PCA analysis.

The main trait used by Hershkovitz (1987) and Marsh (2014) to distinguish these two species was the colour of the face hairs of adult males (and also females by Marsh 2014). Yet, the considerable variability in this trait across their range in the Guiana Shield suggests it may be an unreliable diagnostic feature (Cleuton Miranda, pers. obs.). Notably, Kühl’s (1820) type specimens of *P. rufibarbata* (juvenile) and *P. ochrocephala* (subadult) in the Leiden Museum, from Suriname and French Guiana, respectively, were considered junior synonyms of *P. pithecia* by Hershkovitz (1987, see also Vieira, 1955; Hill, 1960; Napier, 1976; Marsh 2014) based on their localities. They were named for their reddish-brown faces—a red beard and an ochre head, respectively. A closer examination of the geographic distribution of this trait is needed to determine its usefulness as a diagnostic character and the validity of *P. pithecia* and *P. chrysocephala* as distinct species.

Our molecular analysis supports the four members of Hershkovitz’s (1987) *monachus* group. *Pithecia albicans* is morphologically the most distinctive in the genus due to its larger size and striking orange-to-blonde pelage, even though Cabrera (1957), Hill (1960) and Napier (1976) considered it to be a subspecies of *monachus.* Cruz Lima (1945), Hershkovitz (1979, 1987), Groves (2001, 2005) and Marsh (2014) placed it as a distinct species. In our analysis, *Pithecia albicans* is the basal species in this group, as was indicated by Marroig and Cheverud (2004). It has a restricted distribution in the lower Purus-Juruá interfluvium (Fig. 1).

Our Bayesian analysis using the cyt *b* dataset showed some geographical structuring with samples from the west of the Rio Juruá and west of the Rio Japurá forming a low-supported clade. This clade corresponds to Hershkovitz’s (1987) *P. monachus.* Hershkovitz accepted only *milleri* from the upper Río Caquetá basin, west of the Río Caguán south to the upper Putumayo, as a subspecies, as did Hill (1960) and Groves (2001), with *hirsuta* and *napensis* as junior synonyms. Marsh’s (2014) assessment placed all these sakis as species and added *P. cazuzai*. For the genomic analysis, we had only four samples, all within the geographical distribution of *P. hirsuta* (*sensu* Marsh 2014), which formed a well-supported clade of their own. Considering the limitations of our dataset, more samples are needed to determine if Hershkovitz’s (1987) consideration of *P. hirsuta*, *P. inusta*, and *P. napensis* as junior synonyms of *P. monachus* can be supported, or if Marsh’s recognition of these as separate species, along with *P. cazuzai* is valid. However, at present, our cyt *b* results indicate we recognise *P. monachus*, but with *P. hirsuta*, *P. inusta*, and *P. cazuzai*, as well as *P. m. milleri* as junior synonyms.

In our cyt *b* Bayesian analysis, samples from east of the Rio Purus and the west bank of the Rio Tapajós form a well-supported separate clade, except for specimens MPEG 752 and MPEG 851, which grouped with those from the west of the Rio Juruá. These two specimens, assigned to *P. irrorata*, are from the state of Acre in Brazil, located on the southern edge of this species’ distribution and at the headwaters of the rios Purus and Juruá, may have introgressed with neighbouring *P. monachus* or *P. vanzolinii*.

This clade delimited by the Rio Purus and Rio Tapajos corresponds to Hershkovitz’s (1987) *P. irrorata*, including the subspecies *vanzolinii*, and to Marsh’s (2014) *P. irrorata*, along with *P. mittermeieri, P. rylandsi, P. pissinattii*, and *P. vanzolinii.* It also includes the Rio Jamari sakis, collected in 1988 by H. Schneider during the construction of the Samuel Hydroelectric Dam, east of the Rio Madeira in the state of Rondônia. Marsh (2014; Appendix II) found that the Jamari females were unlike any others in the genus, but that the males were variably a mix with features of *P. irrorata*, *P. mittermeieri* and *P. rylandsi.* The ddRAD genomic analysis retrieved only one lineage in this clade. The PCA analysis also showed a single cluster including all these samples. Notably, 32% of the genetic variation is also explained by PC1 and PC3 and shows some structuring (Fig. 5C) between samples found in the Purus-Madeira interfluvium (*P. pissinatti*) and samples from the Tapajós-Madeira interfluvium (*P. mittermeieri*).

Gray (1842) mentioned *P. irroratus* [sic] with no description or reference to a type specimen. As such, it is a *nomen nudum*, as explained by Serrano-Villavicencio et al. (2019). Gray (1843a, p.14) provided the first description and an illustration but no type locality. Gray (1843a, 1843b) indicated that it had been collected during the voyage of the ship H.M.S. Sulphur, which stopped off in Rio de Janeiro. There is no record of any incursions by the expedition into Brazilian Amazonia. Marsh (2014) suggested that it might have been collected during the ship’s visit to Lima, but that seems unlikely. Hershkovitz (1987) imagined that it was taken from a market and arbitrarily restricted the type locality to the left bank of the Rio Tapajós in the Amazônia National Park, Amazonas, Brazil. The holotype is based on the illustration of *Pithecia irrorata* in Gray (1843a) (reproduced in Marsh (2014) and Serrano-Villavicencio et al. (2019)). The holotype today is restricted to a skull, accession number 101a, in the British Museum of Natural History; the skin has been lost (Napier, 1976).

Although it is impossible to determine the provenance of Gray’s (1843a) *irrorata*, our genomic results show that individuals from the east bank of the Rio Purus and the left bank of the Rio Tapajós are a single species, which Hershkovitz (1987) ascribed to *P. irrorata*, which supports previous findings based on the pattern of pelage coloration (Serrano-Villavicencio et al., 2019) and baldness of the face which is unique in this species.

In our cyt *b* Bayesian analysis, three samples of *P. vanzolini* grouped with samples from the west of the Rio Juruá. However, our ddRAD genomic analysis positioned our single sample of *P. vanzolinii* as a sister taxon to *P. mittermeieri + P. pissinattii*. This finding aligns with Hershkovitz’s (1987) recognition of the affinity between *P. vanzolinii* and his *P. irrorata*, and it also supports Marsh’s (2014) classification of *P. vanzolinii* as a distinct species. The geographic proximity of *P. vanzolinii* to sakis west of the Rio Juruá may account for some introgression across species boundaries in the headwaters of the Rio Juruá. This potential gene flow between geographically close populations could explain the observed grouping in the cyt *b* analysis, despite the genomic evidence supporting the distinctiveness of *P. vanzolinii*.

The cyt *b* Bayesian analysis identified, albeit with low support, a separate clade containing samples from Ecuador and within the ranges of *P. aequatorialis* and *P. isabela,* and a single sample of *P. inusta* (FMNH64270) from Peru (Fig 3). We did not have genomic data to confirm this separation, so fresh samples of these species will be necessary to determine if Hershkovitz’s (1987) *P. aequatorialis* is a distinct species, possibly with *P. isabela* as its junior synonym. Finally, two samples of *P. monachus* (MPEG 1831, FMNH 88862) are sisters to the clade containing all samples minus *P. albicans*. This result is somewhat unexpected because these samples do not group with the clade that includes samples from the Sedimentary Basin and Brazilian Shield regions. It is possible that low genetic diversity in the total alignment and potential issues with base calling due to poor DNA amplification might contribute to this unexpected phylogenetic placement.

The distinctive *Pithecia napensis* was considered a synonym of *P. monachus* by Cabrera (1957), Hill (1960), Napier (1976), and Hershkovitz (1987), and a phylogenetic study has still to be carried out to confirm or otherwise the proposal of Marsh (2014). In appearance and geographic distribution, it would seems to be aligned with *P. aequatorialis* and *P. isabela* and, in the dullest manifestation of a hypothetical clade, with *P. inusta.* We also failed to obtain samples for *Pithecia rylandsi*. Serrano-Villavicencio et al. (2019) argued that it is a synonym of *P. irrorata*, along with *P. mittermeieri* and *P. pissinnattii.* Rylands et al. (2024) considered that it was probably not a valid taxon, based not only on the analysis of Serrano-Villavicencio et al. (2019) but also because its geographic distribution, as proposed by Marsh (2014), indicated that it would be sympatric with *P. irrorata* in the west of its range (south-east Peru and northern Bolivia), and even with *P. mittermeieri* in the east of its range in Rondônia, which in ecological and behavioural terms they felt was highly unlikely.

The split between *Pithecia* and its sister clades (uakaris and bearded sakis) occurred during the Miocene (∼13 million years ago) (Kuderna et al., 2023). The first split in *Pithecia* is more recent, dating to around 3 million years ago, coinciding with the formation of the Amazon transcontinental drainage (Campbell et al., 2006; Latrubesse et al., 2010; Rossetti et al., 2015). The ancestral *Pithecia* population likely had a wide distribution, which was severed by the formation of the Rio Amazonas. Subsequent diversification within *Pithecia* occurred during the late Pleistocene, suggesting rapid geographic expansion and diversification during the Quaternary.

Despite its wide geographic distribution, *Pithecia* shows low genetic diversity. Large rivers in the Amazon typically drive speciation (Boubli et al., 2015; Ribas et al., 2012), yet some *Pithecia* lineages span across major rivers such as the Solimões, Madeira, and potentially the Tapajós. The relatively numerous lineages are, however, consistent with those seen in other pithecine genera: five species in *Chiropotes* and eight in *Cacajao* (Boubli et al., 2008; Carneiro et al., 2023; Silva et al., 2022, 2024).

Our molecular results have shown an unexpectedly low genetic diversity in *Pithecia* mitochondrial DNA. Our limited genomic data offered more phylogenetic resolution, indicating the existence of at least six distinct species: *P. pithecia*, *P. chrysocephala*, *P. albicans*, *P. irrorata*, *P. monachus*, and *P. vanzolinii.* However, the striking morphological distinctions and geographic structuring reported by Marsh (2014) (and Hershkovitz 1987) led this author to recognise 16 taxa under the Phylogenetic Species Concept (PSC, *sensu* Cracraft 1989). Given her findings, a more comprehensive genomic examination, using a larger and more representative sampling, is necessary to test her hypothesis with molecular data.

## Competing interests

The Authors declare no financial or personal conflict of interest.

## Author contributions

**J.P.B**. conceived and designed the study, acquired funding and organised sample acquisition, interpreted the findings, and led the writing of the manuscript. **F.E.S.** provided samples, led genomic analyses, interpreted findings, participated in writing the manuscript, and created figures with input from **J.P.B**.. **P.S.B., W.F., R.B.** extracted and sequenced the samples, led the phylogenetic and species delimitation analyses, created figures and contributed to the writing of the manuscript. **R.C.A., C.M., R.R., M.G., J.V., M.M.**, provided samples and contributed to the writing of the manuscript, **P.B.** curated the sample locations, produced Figure 1 and edited the manuscript, **M.N.F.S**. conceived the study, interpreted findings, and provided samples. **A.B.R**. and **R.M**. conceived the study, interpreted findings, and participated in writing the manuscript. **A.H.H.** provided samples from the Spix syntypes and edited the manuscript. **C.R.** led the extraction and sequencing of mitogenomes of Spix syntypes, interpreted the findings, and participated in the writing of the manuscript. **T.H.** and **I.P.F**. conceived and designed the study, acquired funding and organised sample acquisition, interpreted the findings, extracted and sequenced samples, and participated in writing the manuscript. All authors read and approved the final version of the manuscript.

## Supporting information

https://docs.google.com/document/d/1EOBf5sOnrBZLwdRBtaSMO-GD-hkp4TR2/edit?usp=drive_link&ouid=114756968278581238731&rtpof=true&sd=true

## Acknowledgements

JPB was supported by a grant from NERC (NE/T000341/1). Molecular analyses and field expeditions were funded by CNPq/ SISBIOTA-BioPHAM (563348/2010) to IPF, CAPES/PRO-AMAZONIA/ AUXPE (3261/2013) to IPF and HS), NSF/FAPESP “Dimensions of Biodiversity” (NSF1241066 and FAPESP12/50260-6) to Joel Cracraft and Lucia Lohman, the Primate Action Fund of the Margot Marsh Biodiversity Foundation (6002856, granted to RCA and TH), and ARPA/ICMBio (granted to RCA). F.E.S. gratefully acknowledges the financial support from the European Union’s Horizon 2020 research and innovation programme under the Marie Skłodowska-Curie grant agreement (801505), the Fonds National de la Recherche Scientifique (F.R.S.-FNRS, Belgium; grant 40017464) Brazilian National Council for Scientific and Technological Development (CNPq) (Processes 303286/2014-8, 303579/2014-5, 200502/2015-8, 302140/2020-4, 300365/2021-7, 301407/2021-5, #301925/2021-6) and the Gordon and Betty Moore Foundation (Grant 5344 to the Mamirauá Institute). Permission to conduct fieldwork and to collect tissue samples was granted by IBAMA and ICMBio (license N° 005/2005 – CGFAU/LIC and ICMBio 32095-1). RCA receives a doctorate scholarship (CNPq140039/2018-0) and is also grateful to the managers and staff of Thaimaçu Lodge, to DEMUC/SEMA offices in Amazonas State and to the team of ICMBio bureau in Itaituba, Pará State, for supporting fieldwork in southern Amazonia. We are grateful to Consórcio Hidrelétrico Teles Pires and Biota–Projetos e Consultoria Ambiental. We are grateful to Conservation International Brazil, Sema Pará and Museu Paraense Emílio Goeldi for the collection of the CN specimens of *P. chrysocephala* and *P. pithecia* from the Guiana shield (RR and CM). We are also grateful to Ingrid Macedo for producing the photos of the museum specimens (INPA). The collection of RR specimens from Mato Grosso state was supported by the Fundação de Amparo à Pesquisa do Estado de Mato Grosso (FAPEMAT, process #477017/2011).

## Notes

### Competing Interest Statement

The authors have declared no competing interest.

