## Supplementary material for "First Comprehensive Examination of the Molecular Phylogenetics of Saki Monkeys, Genus *Pithecia* Desmarest, 1804, Reveals an Unexpectedly Low Taxonomic Diversity": https://docs.google.com/document/d/1EOBf5sOnrBZLwdRBtaSMO-GD-hkp4TR2/edit?usp=drive_link&ouid=114756968278581238731&rtpof=true&sd=true

Tabela 1 -  Primers used
in Figueiredo (2005)

|  |  |  |
| --- | --- | --- |
| Primer | Direction | Sequence (5’ → 3’) |
| L-14727 | Forward | TGATATGAAAAACCATCGTTG |
| Lulu-125 | Forward | TCATCCAAATCATCACAGGCC |
| Ana-208 | Forward | ACCCGAGATGTAAACTACGG |
| Jeff-353 | Forward | TTTTATTACTTACAACTATAGC |
| Tom-475 | Forward | GACCTAGTACAATGAATCTGAGG |
| Pepe-592 | Forward | CTGTTTCTGCATGACACAGG |
| Terry-712 | Forward | CTTCTAATAAGCCTAACCC |
| Jen-827 | Forward | TTGCATACGCAATTCTACGATC |
| Rachel-1034 | Forward | ACCCTTTTATTAGCATTGGCC |
| Tyana-154 | Reverse | GAGTCTGATGTGTAATGTATAGC |
| Hafa-263 | Reverse | AATAGACATATGAAGAATAGGG |
| Alex-378 | Reverse | GAACATAGCCTATAAATGCTG |
| Gina-538 | Reverse | AGGCAAGATAAAGTGGAAGG |
| Michi-622 | Reverse | CGGATGTCAGTCCTGATGG |
| Jutta-757 | Reverse | AGCTGGGGTGTAGTTGTCTGG |
| Trina-862 | Reverse | AGGGCTAAGACGCCTCCTAG |
| Xhosa-1060 | Reverse | TAAGAAAGTATAAGGTGGATGC |
| H-15915 | Reverse | TCATCTCCGGTTTACAAGAC |
